# 4DNvestigator: Time Series Genomic Data Analysis Toolbox

**DOI:** 10.1101/2020.01.08.898387

**Authors:** Stephen Lindsly, Can Chen, Sijia Liu, Scott Ronquist, Samuel Dilworth, Michael Perlman, Indika Rajapakse

## Abstract

Data on genome organization and output over time, or the 4D Nucleome (4DN), require synthesis for meaningful interpretation. Development of tools for the efficient integration of these data is needed, especially for the time dimension. We present the “4DNvestigator”, a user-friendly network based toolbox for the analysis of time series genome-wide genome structure (Hi-C) and gene expression (RNA-seq) data. Additionally, we provide methods to quantify network entropy, tensor entropy, and statistically significant changes in time series Hi-C data at different genomic scales.

**Availability:** https://github.com/lindsly/4DNvestigator

## 1 Introduction

4D nuclear organization (4D Nucleome, 4DN) is defined by the dynamical interaction between 3D genome structure and function [1, 2, 3]. To analyze the 4DN, genome-wide chromosome conformation capture (Hi-C) and RNA sequencing (RNA-seq) are often used to observe genome structure and function, respectively (Figure 1A). The availability and volume of Hi-C and RNA-seq data is expected to increase as high throughput sequencing costs decline, thus the development of methods to analyze these data is imperative. The relationship of genome structure and function has been studied previously [4, 5, 6, 3, 7], yet comprehensive and accessible tools for 4DN analysis are underdeveloped. The 4DNvestigator is a unified toolbox that loads time series Hi-C and RNA-seq data, extracts important structural and functional features (Figure 1B), and conducts both established and novel 4DN data analysis methods. We show that network centrality can be integrated with gene expression to elucidate structural and functional changes through time, and provide relevant links to the NCBI and GeneCards databases for biological interpretation of these changes [8, 9]. Furthermore, we utilize entropy to quantify the uncertainty of genome structure, and present a simple statistical method for comparing two or more Hi-C matrices.

**Figure 1:**
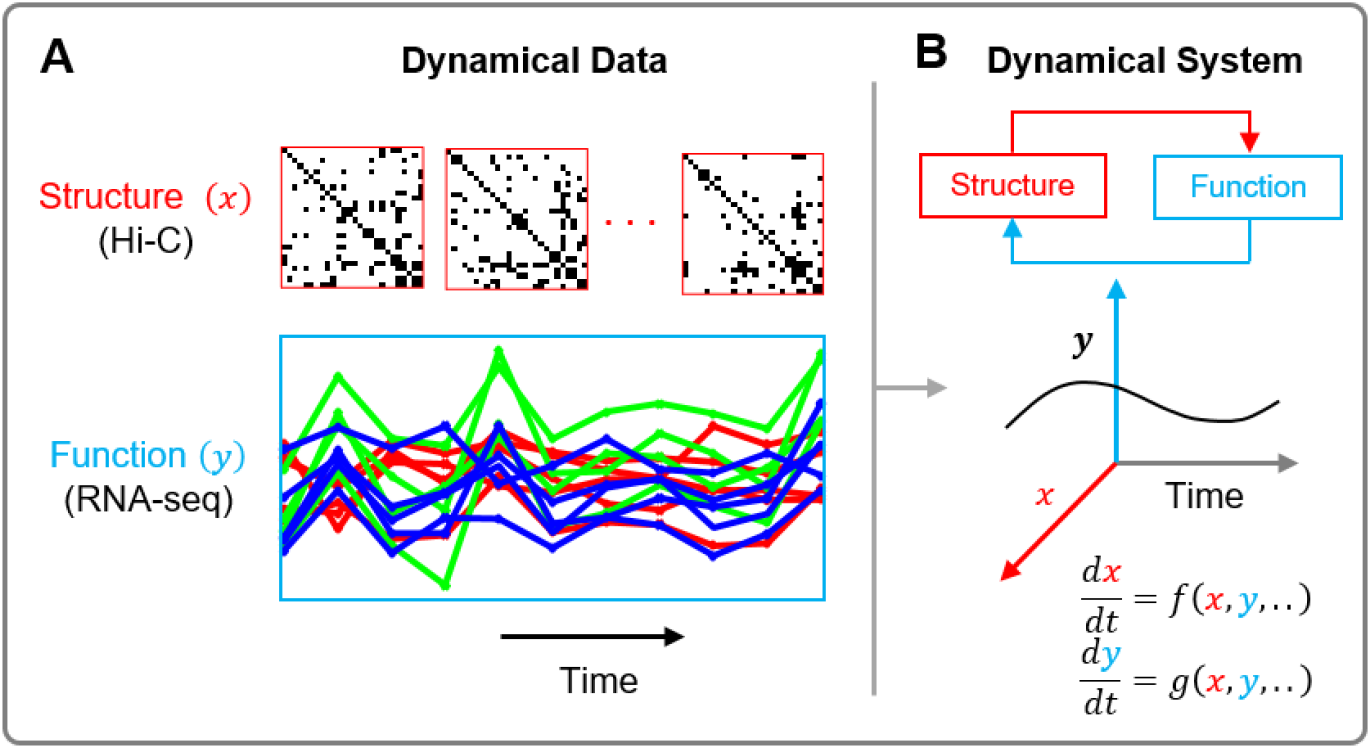
The 4D Nucleome. (A) Representative time series Hi-C and RNA-seq data correspond to genome structure and function, respectively. (B) Genome structure and function are intimately related. The 4DNvestigator integrates and visualizes time series data to study their dynamical relationship.

## 2 Materials and Methods

An overview of the 4DNvestigator workflow is depicted in Figure 2, and a Getting Started document is provided to guide the user through the main functionalities of the 4DNvestigator. The 4DNvestigator takes processed Hi-C and RNA-seq data as input, along with a metadata file which describes the sample and time point for each input Hi-C and RNA-seq file (See Supplementary Materials “Data Preparation”). A number of novel methods for analyzing 4DN data are included within the 4DNvestigator and are described below.

**Figure 2:**
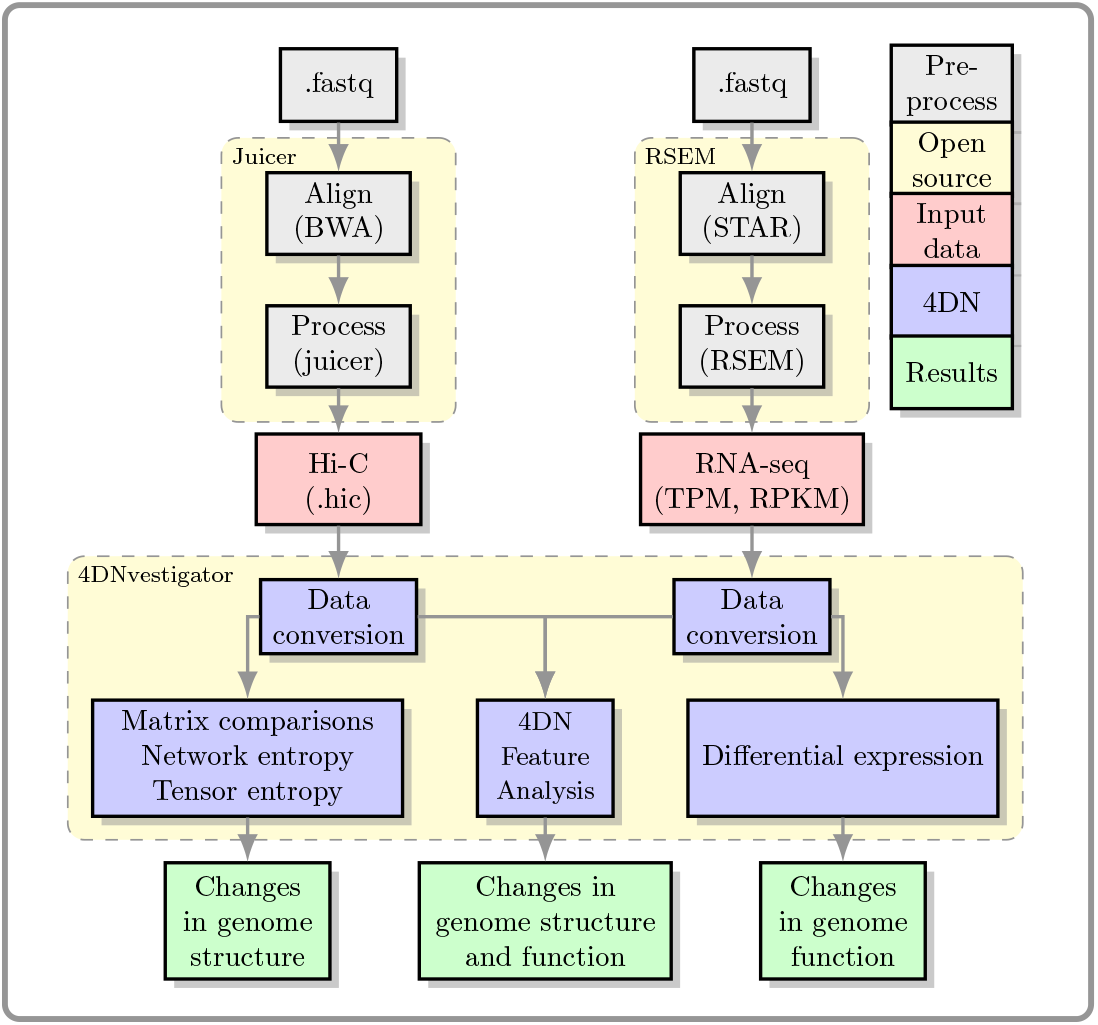
Overview of the 4DNvestigator data processing pipeline. Within this diagram, 4DN refers to the 4DNvestigator.

### 2.1 4DN Feature Analyzer

The “4DN feature analyzer” quantifies and visualizes how much a genomic region changes in structure and function over time. To analyze both structural and functional data, we consider the genome as a network. Nodes within this network are genomic loci, where a locus can be a gene or a genomic region at a particular resolution (i.e. 100 kb or 1 Mb bins). Edges in the genomic network are the relationships or interactions between genomic loci.

**Algorithm 1:**
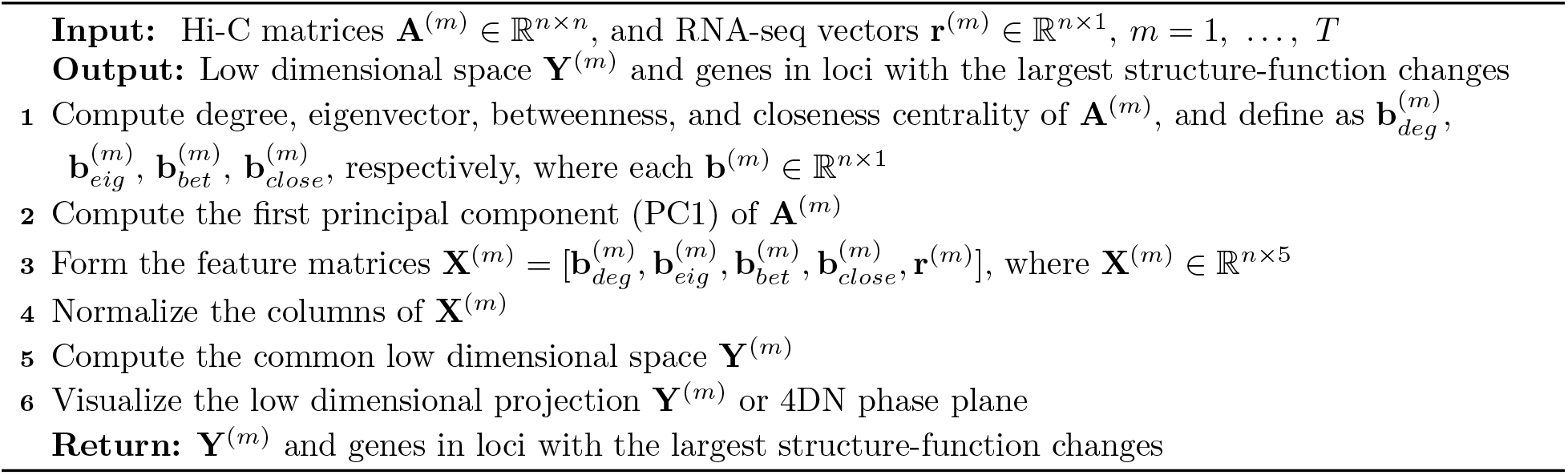
4DN feature analyzer

#### Structural Data

Structure in the 4DN feature analyzer is derived from Hi-C data. Hi-C determines the edge weights in our genomic network through the frequency of contacts between genomic loci. To analyze genomic networks, we adopt an important concept from network theory called centrality. Network centrality is motivated by the identification of nodes which are the most “central” or “important” within a network [10]. The 4DN feature analyzer uses *degree*, *eigenvector*, *betweenness*, and *closeness* centrality (step 1 of Algorithm 1), which have been shown to be biologically relevant [7]. For example, eigenvector centrality can identify structurally defined regions of active/inactive gene expression, since it encodes clustering information of a network [7, 11]. Additionally, betweenness centrality measures the importance of nodes in regard to the flow of information between pairs of nodes. Boundaries between euchromatin and heterochromatin, which often change in reprogramming experiments, can be identified in a genomic network through betweenness centrality [7].

#### Functional Data

Function in the 4DN feature analyzer is derived from gene expression through RNA-seq. Function is defined as the log2 transformation of Transcripts Per Million (TPM) or Reads Per Kilobase Million (RPKM). For regions containing more than one gene, the mean expression of all genes within the region is used.

#### Integration of Data

Hi-C data is naturally represented as a matrix of contacts between genomic loci. Network centrality measures are one-dimensional vectors that describe important structural features of the genomic network. We combine network centrality with RNA-seq expression to form a structure-function “feature” matrix that defines the state of each genomic region at each time point (Figure 3A, step 3 of Algorithm 1). Within this matrix, rows represent genomic loci and columns are the centrality measures (structure) and gene expression (function) of each locus. The z-score for each column is computed to normalize the data (step 4 of Algorithm 1).

**Figure 3:**
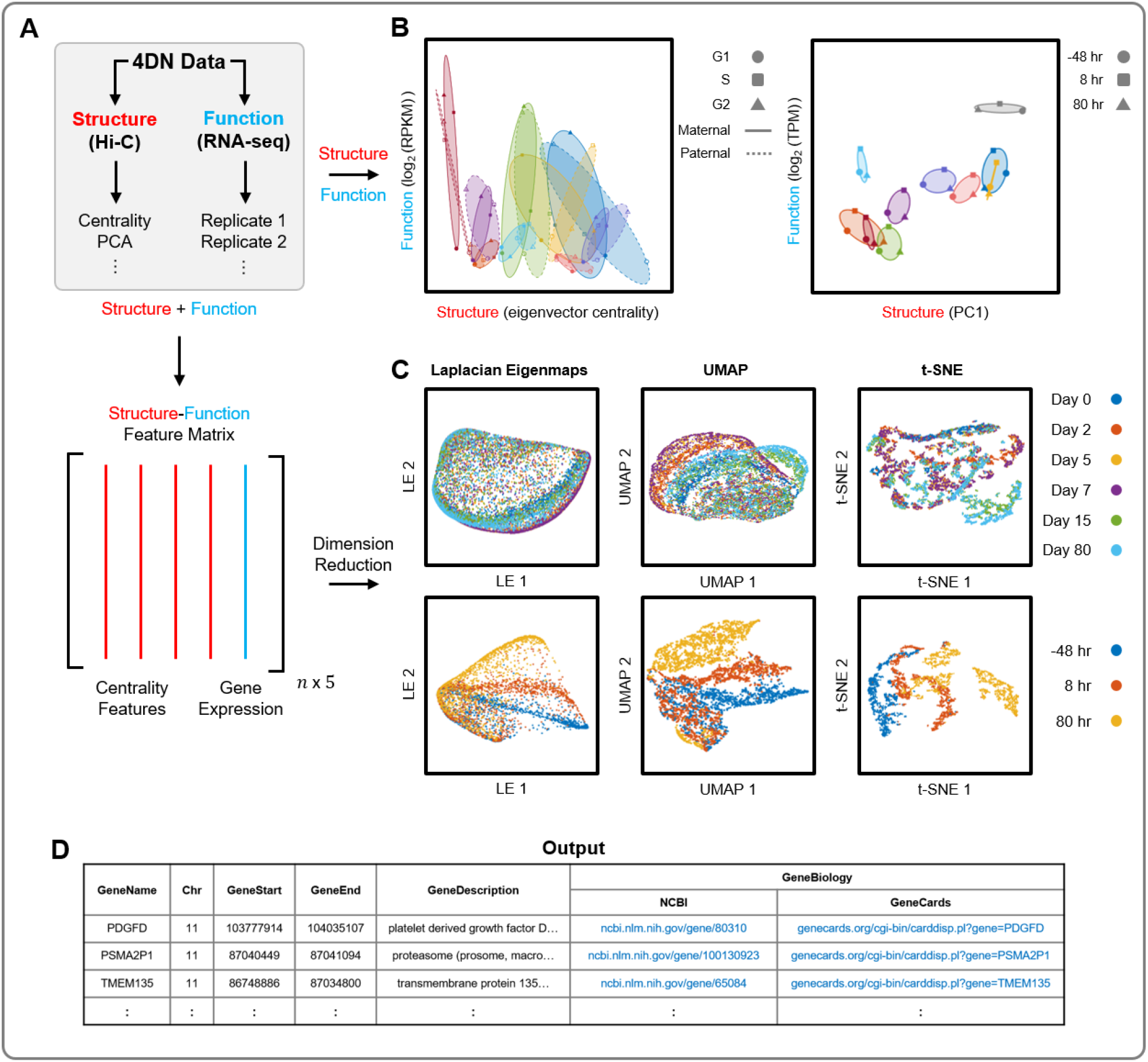
4DN feature analyzer. (A) 4DN data is input to the 4DN feature analyzer. Top: Structure data (Hi-C) is described using one dimensional features for compatibility with function data (RNA-seq). Bottom: Multiple structural features and function data are integrated into the structure-function feature matrix. (B) The 4DN feature analyzer can use structure and function data directly to visualize a system’s dynamics using a phase plane. Structure defines the *x*-axis (left: eigenvector centrality, right: PC1) and function defines the *y*-axis (left: log2(RPKM), right: log2(TPM)), and points show structure-function coordinates through time. Left: Maternal and paternal alleles of nine cell cycle genes through G1, S, and G2/M phases of the cell cycle (adapted from [15]). Right: Top ten genomic regions (100 kb) with the largest changes in structure and function during cellular reprogramming [7]. (C) Multiple dimension reduction techniques can be used to visualize the 4DN feature analyzer’s structure-function feature matrix (from left to right: LE, UMAP, and t-SNE). Top: 100 kb regions of Chromosome 4 across six time points during cellular differentiation [28]. Bottom: 100 kb regions of Chromosome 11 across three time points during cellular reprogramming [7]. (D) Example output of the 4DN feature analyzer. The output includes genes contained in loci with the largest changes, and links to their NCBI and GeneCards database entries [8, 9].

#### 4DN Analysis

The 4DN feature analyzer reduces the dimension of the structure-function feature matrix for visualization and further analysis (steps 5 and 6 of Algorithm 1). We include the main linear dimension reduction method, Principal Component Analysis (PCA), and multiple nonlinear dimension reduction methods: Laplacian Eigenmaps (LE) [12], t-distributed Stochastic Neighbor Embedding (t-SNE) [13], and Uniform Manifold Approximation and Projection (UMAP) [14] (Figure 3C). These methods are described in more detail in Supplementary Materials “Dimension Reduction”. The 4DN feature analyzer can also visualize the dynamics of genome structure and function using the 4DN phase plane (step 6 of Algorithm 1) [3, 15]. We designate one axis of the 4DN phase plane as a measure of genome structure (e.g. eigenvector centrality) and the other as a measure of genome function (gene expression). Each point on the phase plane represents the structure and function of a genomic locus at a specific point in time (Figure 3B). The 4DN feature analyzer identifies genomic regions and genes with large changes in structure and function over time, and provides relevant links to the NCBI and GeneCard databases [8, 9].

### Additional 4DNvestigator Tools

#### General Structure and Function Analysis

The 4DNvestigator also includes a suite of previously developed Hi-C and RNA-seq analysis methods. Euchromatin and heterochromatin compartments can be identified from Hi-C [4, 16], and regions that change compartments between samples are automatically identified. Significant changes in gene expression between RNA-seq samples can be determined through differential expression analysis using established methods [17].

#### Network Entropy

Entropy measures the amount of uncertainty within a system [18]. We use entropy to quantify the organization of chromatin structure from Hi-C data, where higher entropy corresponds to less structural organization. Since Hi-C is a multivariate analysis measurement (each contact coincidence involves two variables, the two genomic loci), we use multivariate entropy as follows:

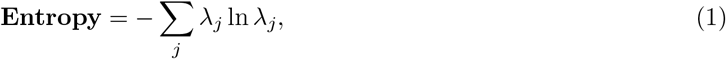

where *λ_i_* represents the dominant features of the Hi-C contact matrix. In mathematics, these dominant features are called eigenvalues [19]. Biologically, genomic regions with high entropy likely correlate with high proportions of euchromatin, as euchromatin is more structurally permissive than heterochromatin [20, 21]. Furthermore, entropy can be used to quantify stemness, since cells with high pluripotency are less defined in their chromatin structure [22]. We provide the full algorithm for network entropy and calculate the entropy of Hi-C data from multiple cell types in Supplementary Materials “Network Entropy”.

#### Tensor Entropy

The notion of transcription factories supports the existence of simultaneous interactions involving three or more genomic loci [23]. This implies that the configuration of the human genome can be more accurately represented by *k*-uniform hypergraphs, a generalization of networks in which each edge can join exactly *k* nodes (e.g. a standard network is a 2-uniform hypergraph). We can construct *k*-uniform hypergraphs from Hi-C contact matrices by computing the multi-correlations of genomic loci. Tensor entropy, an extension of network entropy, measures the uncertainty or disorganization of uniform hypergraphs [24]. Tensor entropy can be computed from the same entropy formula (1) with generalized singular values *λ_j_* from tensor theory [24, 25]. We provide the definitions for multi-correlation and generalized singular values, the algorithm to compute tensor entropy, and an application of tensor entropy on Hi-C data in Supplementary Materials “Tensor Entropy”.

#### Larntz-Perlman Procedure

The 4DNvestigator includes a statistical test, proposed by Larntz and Perlman (the LP procedure), that compares correlation matrices [26, 27]. The LP procedure is applied to correlation matrices from Hi-C data, and is able to determine whether multiple Hi-C samples are significantly different from one another. Suppose that **C**^(*m*)^ ∈ ℝ*n*×*n* are the sample correlation matrices of Hi-C contacts with corresponding population correlation matrices **P**^(*m*)^ ∈ ℝ^*n*×*n*^ for *m* = 1, 2,…, *k*. The null hypothesis is *H*_0_: **P**^(1)^ =⋯= **P**^(*k*)^. First, compute the Fisher z-transformation **Z**^(*m*)^ by

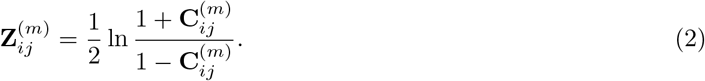

Then form the matrices **S**^(*m*)^ such that

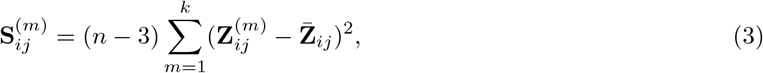

where, 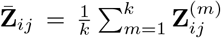. The test statistic is given by *T* = max_*ij*_ **S**_*ij*_, and *H*_0_ is rejected at level *α* if 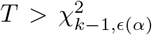 where 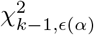 is the chi-square distribution with *k* − 1 degree of freedom, and *ϵ*(*α*) = (1 − *α*)^2/(*n*(*n*−1))^ is the Šidák correction. Finally, calculate the *p*-value at which 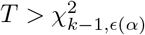. We note that this *p*-value is conservative, and that the the actual *p*-value may be smaller depending upon the amount of correlation among the variables. The LP procedure determines the statistical significance of any differences between multiple Hi-C samples for a genomic region of interest. We provide benchmark results of the LP procedure with other Hi-C comparison methods in Supplementary Materials “LP Procedure for Comparing Hi-C Matrices”.

## 3 Results

We demonstrate how the 4DN feature analyzer can process time series structure and function data (Figure 3A) with three examples (Figure 3B-D).

### Example 1: Cellular Proliferation

Hi-C and RNA-seq data from B-lymphoblastoid cells (NA12878) capture the G1, S, and G2/M phases of the cell cycle for the maternal and paternal genomes [15]. We visualize the structure-function dynamics of the maternal and paternal alleles for nine cell cycle regulating genes using a phase plane (Figure 3B, left). We are interested in the importance of these genes within the genomic network through the cell cycle, so we use eigenvector centrality as the structural measure. This analysis highlights the coordination between the maternal and paternal alleles of these genes through the cell cycle.

### Example 2: Cellular Differentiation

We constructed a structure-function feature matrix from time series Hi-C and RNA-seq data obtained from differentiating human stem cells [28]. These data consist of six time points which include human embryonic stem cells, mesodermal cells, cardiac mesodermal cells, cardiac progenitors, primitive cardiomyocytes, and ventricular cardiomyocytes [28]. We analyze Chromosome 4 across the six time points in 100 kb resolution by applying three dimension reduction techniques to the structure-function feature matrix: LE, UMAP, and t-SNE (Figure 3C, top). There is a better separation of the cell types during differentiation using UMAP and t-SNE than from LE. The optimal methods for visualization and analysis are data dependent, so the 4DNvestigator offers multiple tools for the user’s own exploration of their data.

### Example 3: Cellular Reprogramming

Time series Hi-C and RNA-seq data were obtained from an experiment that reprogrammed human dermal fibroblasts to the skeletal muscle lineage [7]. We analyze samples collected 48 hr prior to, 8 hr after, and 80 hr after the addition of the transcription factor MYOD1. The ten 100 kb regions from Chromosome 11 that varied most in structure and function are visualized using a phase plane in Figure 3B (right). We also construct a structure-function feature matrix of Chromosome 11 in 100 kb resolution. Similar to the differentiation data analysis, we use LE, UMAP, and t-SNE to visualize the structure-function dynamics. These low dimensional projections show the separation of the three time points corresponding to before, during, and after cellular reprogramming (Figure 3C, bottom). We show an example output of the 4DN feature analyzer, which highlights genes contained in the genomic loci that have the largest structure-function changes through time and provides links to the NCBI and GeneCards database entries for these genes (Figure 3D) [8, 9].

## 4 Discussion

The 4DNvestigator provides rigorous and automated analysis of Hi-C and RNA-seq time series data by drawing on network theory, information theory, and multivariate statistics. It also introduces a simple statistical method for comparing Hi-C matrices, the LP procedure. The LP procedure is distinct from established Hi-C matrix comparison methods, as it takes a statistical approach to test for matrix equality, and allows for the comparison of many matrices simultaneously. Thus, the 4DNvestigator provides a comprehensive toolbox that can be applied to time series Hi-C and RNA-seq data simultaneously or independently. These methods are important for producing rigorous quantitative results in 4DN research.

## 5 Acknowledgments

We would like to thank Dr. Thomas Ried, Charles Ryan, and Gabrielle Dotson for feedback on the manuscript and helpful discussions.

## 8 Supplemental Material

Please refer to the file ‘4DNvestigator Supplementary Materials’ for additional details on the 4DNvestigator’s installation process, methods, and data availability.

## References

[1] Job Dekker, Andrew S. Belmont, Mitchell Guttman, Victor O. Leshyk, John T. Lis, Stavros Lomvardas, Leonid A. Mirny, Clodagh C. O’Shea, Peter J. Park, Bing Ren, Joan C. Ritland Politz, Jay Shendure, and Sheng Zhong. The 4D nucleome project. Nature, 549(7671):219–226, 2017.

[2] Thomas Ried and Indika Rajapakse. The 4D Nucleome. Methods, 123:1–2, 2017.

[3] Haiming Chen, Jie Chen, Lindsey a. Muir, Scott Ronquist, Walter Meixner, Mats Ljungman, Thomas Ried, Stephen Smale, and Indika Rajapakse. Functional organization of the human 4D Nucleome. Proceedings of the National Academy of Sciences, 112(26):8002–8007, 2015.

[4] Erez Lieberman-aiden, Nynke L Van Berkum, Louise Williams, Maxim Imakaev, Tobias Ragoczy, Agnes Telling, Ido Amit, Bryan R Lajoie, Peter J Sabo, Michael O Dorschner, Richard Sandstrom, Bradley Bernstein, M A Bender, Mark Groudine, Andreas Gnirke, John Stamatoyannopoulos, Leonid A Mirny, Eric S. Lander, and Job Dekker. Comprehensive Mapping of Long-Range Interactions Reveals Folding Principles of the Human Genome. Science, 326:289–294, 2009.

[5] Jesse R. Dixon, Inkyung Jung, Siddarth Selvaraj, Yin Shen, Jessica E. Antosiewicz-Bourget, Ah Young Lee, Zhen Ye, Audrey Kim, Nisha Rajagopal, Wei Xie, Yarui Diao, Jing Liang, Huimin Zhao, Victor V. Lobanenkov, Joseph R. Ecker, James A. Thomson, and Bing Ren. Chromatin architecture reorganization during stem cell differentiation. Nature, 518(7539):331–336, 2 2015.

[6] Jesse R. Dixon, Siddarth Selvaraj, Feng Yue, Audrey Kim, Yan Li, Yin Shen, Ming Hu, Jun S. Liu, and Bing Ren. Topological domains in mammalian genomes identified by analysis of chromatin interactions. Nature, 485(7398):376–380, 2012.

[7] Sijia Liu, Haiming Chen, Scott Ronquist, Laura Seaman, Nicholas Ceglia, Walter Meixner, Pin-Yu Chen, Gerald Higgins, Pierre Baldi, Steve Smale, Alfred Hero, Lindsey A. Muir, and Indika Rajapakse. Genome Architecture Mediates Transcriptional Control of Human Myogenic Reprogramming. iScience, 6:232–246, 2018.

[8] David L Wheeler, Tanya Barrett, Dennis A Benson, Stephen H Bryant, Kathi Canese, Vyacheslav Chetvernin, Deanna M Church, Michael DiCuccio, Ron Edgar, Scott Federhen, et al. Database resources of the national center for biotechnology information. Nucleic acids research, 36(suppl 1):D13–D21, 2007.

[9] Gil Stelzer, Naomi Rosen, Inbar Plaschkes, Shahar Zimmerman, Michal Twik, Simon Fishilevich, Tsippi Iny Stein, Ron Nudel, Iris Lieder, Yaron Mazor, et al. The genecards suite: from gene data mining to disease genome sequence analyses. Current protocols in bioinformatics, 54(1):1–30, 2016.

[10] Mark Newman. Networks: an introduction. Oxford university press, New York, 2010.

[11] Andrew Y Ng, Michael I Jordan, Yair Weiss, et al. On spectral clustering: Analysis and an algorithm. In NIPS, volume 14, pages 849–856, 2001.

[12] Mikhail Belkin and Partha Niyogi. Laplacian Eigenmaps and Spectral Techniques for Embedding and Clustering. NIPS, 14:585–591, 2001.

[13] Laurens Van Der Maaten and George E Hinton. Visualizing high-dimensional data using t-SNE. Journal of Machine Learning Research, 9:2579–2605, 2008.

[14] Leland McInnes, John Healy, and James Melville. Umap: Uniform manifold approximation and projection for dimension reduction. arXiv preprint arXiv:1802.03426, 2018.

[15] Stephen Lindsly, Wenlong Jia, Haiming Chen, Sijia Liu, Scott Ronquist, Can Chen, Xingzhao Wen, Gabrielle A Dotson, Charles Ryan, Gilbert S Omenn, et al. Functional organization of the maternal and paternal human 4d nucleome. bioRxiv, 2020.

[16] Jie Chen, Alfred Hero, and Indika Rajapakse. Spectral Identification of Topological Domains. Bioinformatics, 32(14):2151–2158, 2016.

[17] Simon Anders and Wolfgang Huber. Differential expression analysis for sequence count data. Genome biology, 11(R106):1–12, 2010.

[18] Thomas M Cover and Joy A Thomas. Elements of information theory. John Wiley & Sons, 2012.

[19] Gilbert Strang. Introduction to Linear Algebra. Cambridge Press, 2016.

[20] Ben D Macarthur and Ihor R Lemischka. Statistical mechanics of pluripotency. Cell, 154(3):484–489, 2013.

[21] I. Rajapakse, M. Groudine, and M. Mesbahi. What can systems theory of networks offer to biology? PLoS computational biology, 8(6):e1002543, 2012.

[22] Eran Meshorer and Tom Misteli. Chromatin in pluripotent embryonic stem cells and differentiation. Nature reviews Molecular cell biology, 7(7):540, 2006.

[23] Peter R Cook and Davide Marenduzzo. Transcription-driven genome organization: a model for chromosome structure and the regulation of gene expression tested through simulations. Nucleic acids research, 46(19):9895–9906, 2018.

[24] C. Chen and I. Rajapakse. Tensor entropy for uniform hypergraphs. IEEE Transactions on Network Science and Engineering, 7(4):2889–2900, 2020.

[25] Lieven De Lathauwer, Bart De Moor, and Joos Vandewalle. A multilinear singular value decomposition. SIAM journal on Matrix Analysis and Applications, 21(4):1253–1278, 2000.

[26] Kinley Larntz and Michael D Perlman. A simple test for the equality of correlation matrices. Rapport technique, Department of Statistics, University of Washington, 141, 1985.

[27] James A Koziol, Joel E Alexander, Lance O Bauer, Samuel Kuperman, Sandra Morzorati, Sean J O’connor, John Rohrbaugh, Bernice Porjesz, Henri Begleiter, and John Polich. A graphical technique for displaying correlation matrices. The American Statistician, 51(4):301–304, 1997.

[28] Yanxiao Zhang, Ting Li, Sebastian Preissl, Maria Luisa Amaral, Jonathan D Grinstein, Elie N Farah, Eugin Destici, Yunjiang Qiu, Rong Hu, Ah Young Lee, et al. Transcriptionally active herv-h retrotransposons demarcate topologically associating domains in human pluripotent stem cells. Nature genetics, 51(9):1380–1388, 2019.

