## Supplementary Materials for "4DNvestigator: Time Series Genomic Data Analysis Toolbox"

### 4DNvestigator: Supplementary Materials

[1]Department of Computational Medicine and Bioinformatics, University of Michigan, Ann Arbor, MI 48109, USA
[2]Department of Mathematics, University of Michigan, Ann Arbor, MI 48109, USA
[3]MIT-IBM Watson AI Lab, IBM Research, Cambridge, MA 02142, USA
[4]Department of Computer Science, Michigan State University, MI 48824, USA
[5]iReprogram, Ann Arbor, MI 48105, USA
[6]Department of Statistics, University of Washington, Seattle, WA 98195, USA

[*]To whom correspondence should be addressed.

February 2, 2021

#### 4DNvestigator Overview and Setup

The 4DNvestigator toolbox can be downloaded and installed from the 4DNvestigator GitHub page. Installation instructions are included in the 4DNvestigator README along with all data used in this manuscript. We provide a script, ExampleScript.m, which shows how to load data into MATLAB and runs each of the four major functionalities of the 4DNvestigator: 4DN feature analyzer, network entropy, tensor entropy, and the Larntz-Perlman procedure. We also provide a Getting Started document which provides a concise description of each of the example functions' parameters and outputs.

##### Data Preparation

The 4DNvestigator requires a metadata table, referred to as an "Index File", to describe the data. Rows correspond to sequencing data samples (RNA-seq or Hi-C), and the columns correspond to data descriptors. The Index File must have 5 columns with the following headers:

- "path" defines the computer path to the sequencing data
- "dataType" defines the type of sequencing data, either "hic" or "rnaseq"
- "sample" defines the sample, (e.g. "treatment" or "control")
- "timePoint" defines the time point of sample, (e.g. 0 or 24)
- "refGenome" defines the reference genome for the sample (e.g. hg19)

Index Files can be .csv, .tsv, or .xls. An example of an Index File is available here.

##### MATLAB Dependencies

The 4DNvestigator requires the following MATLAB Toolboxes:

- Statistics and Machine Learning Toolbox

- Computer Vision Toolbox
- Bioinformatics Toolbox
- Image Processing Toolbox

##### System Requirements

The 4DNvestigator was written and tested on a Windows 10 machine with an Intel Core i7-8700 CPU and 32 Gb RAM using MATLAB R2019b. During testing, we timed each of the core functionalities' example functions using default settings, including figure generation and saving (Supplementary Table S1). We also measured peak RAM usage over function calls and data loading which was approximately 9.5 Gb (higher RAM usage is caused by higher resolution Hi-C matrices). Therefore, we suggest that users run a recent version of MATLAB (2019 or later) on a Windows 10 machine with at least 12 Gb of RAM for optimal performance. Each of the example functions in Supplementary Table S1 can be run with default parameters using ExampleScript.m, and a step-by-step guide of this script can be found in the Getting Started document.

| ExampleScript.m Section | Description | Runtime |
|---|---|---|
| Load RNA-seq and Hi-C Data | Loads RNA-seq and Hi-C data using results from RSEM and Juicer, respectively | ∼240 sec |
| featureAnalyzerExample.m | Example function for the 4DN feature analyzer, called using default settings | ∼30 sec |
| entropyExample.m | Example function for entropy calculation on two cell types, called using default settings | ∼6 sec |
| entropyExampleExpanded.m | Example function for entropy calculation on six cell types, called using default settings | ∼45 sec |
| tensorEntropyExample.m | Example function for tensor entropy calculation on reprogramming data, called using default settings | ∼6 sec |
| lpExample.m | Example function for the Larntz-Perman procedure on three samples, called using default settings | ∼20 sec |

Table S1: Mapping of core 4DNvestigator functionalities and their example functions within the toolbox. Each of these functions are called within ExampleScript.m. Here we list their respective approximate runtimes.

#### Dimension Reduction

The 4DN feature analyzer provides multiple methods for reducing the dimension of structure-function data. The first is Principal Component Analysis (PCA), a linear dimension reduction technique. PCA is performed by computing the eigendecomposition of the centered covariance matrix. PCA uses the eigenvectors associated with the largest eigenvalues as coordinates for the low dimensional space. Unlike PCA, the other dimension reduction techniques included in the 4DN feature analyzer are non-linear. Non-linear methods are based on the idea that high dimensional data often lie on a curved manifold in lower dimension. In the 4DN feature analyzer, we provide the choice between Laplacian Eigenmaps (LE), t-distributed Stochastic Neighbor Embedding (t-SNE), and Uniform Manifold Approximate Projection (UMAP). LE constructs a graph, defined as the similarities between genomic loci in terms of their structural and functional features. It then computes the eigendecomposition of the Laplacian matrix, and uses the eigenvectors associated with the smallest non-zero eigenvalues as the low dimensional coordinates [1]. The Laplacian matrix is calculated by subtracting the graph's adjacency matrix from its degree matrix. The goal of both t-SNE and UMAP is to reduce the dimension of high dimensional data, while preserving the relationships between data. In other words, t-SNE and UMAP optimize the low dimensional mapping of data to ensure that data which are similar to one another in high dimensional space are also close in low dimensional space [2, 3]. UMAP is conceptually similar to t-SNE, but it is better at preserving global structure of data and more computationally efficient [3].

#### Network Entropy

Entropy measures the order within a system, where higher entropy corresponds to more disorder [4]. Here, we apply this measure to Hi-C data to quantify the order in chromatin structure. Biologically, genomic regions with high entropy likely correlate with high proportions of euchromatin, as euchromatin is more structurally permissive than heterochromatin [5, 6]. Furthermore, entropy can be used to quantify stemness, since cells with high pluripotency are less defined in their chromatin structure [7]. Since Hi-C is a multivariate analysis measurement (each contact coincidence involves two variables, the two loci), we use multivariate entropy. The algorithm to compute entropy is given in Algorithm 1. We provide two variants of the algorithm to calculate entropy. The first uses the correlation matrix of $\log_2$ transformed Hi-C data, while the second uses the Laplacian matrix from Hi-C data. We find that the first variant performs better at discriminating the stemness of cell types, so we use this variant as the default for the entropy calculation. For both variants of network entropy, we use the convention $0 \ln 0 = 0$.

**Algorithm 1:** Entropy Computation

**Input:** Hi-C matrix $\mathbf{A} \in \mathbb{R}^{n \times n}$, Entropy variant
**Output:** Entropy

**if** *variant* == *'corr'* **then**
    Compute the correlation matrix $\mathbf{C} = \mathrm{corr}(\log_2(\mathbf{A}))$
    Compute the eigenvalues $\lambda_i$ of $\mathbf{C}$ using eigendecomposition
**else if** *variant* == *'laplacian'* **then**
    Compute the Laplacian matrix $\mathbf{L} = \mathbf{D} - \mathbf{A}$ where $\mathbf{D}$ is the degree matrix of $\mathbf{A}$
    Compute the eigenvalues $\lambda_i$ of $\mathbf{L}$ using eigendecomposition
Normalize the eigenvalues: $\bar{\lambda}_j = \frac{\lambda_j}{\sum_{i=1}^{n} \lambda_i}$
Compute Entropy: $\textbf{Entropy} = -\sum_j \bar{\lambda}_j \ln \bar{\lambda}_j$
**Return:** Entropy

An application of entropy is shown in Figure S1. We use entropy to to quantify stemness on the following samples: human Embryonic Stem Cells (hESC), Human Umbilical Vein Endothelial Cells (HUVEC), human lung fibroblasts (IMR90), Human Foreskin Fibroblast cells (HFFc6), human B-lymphocyte cells (GM12878). These data were obtained from the 4DN Portal and other publicly available datasets [8, 9, 10]. A simple comparison between two cell types can be performed using entropyExample and an example of multiple cell types (Figure S1) can be performed using entropyExampleExpanded. Both examples are called by the script ExampleScript.m.

#### Tensor Entropy

Uniform hypergraphs can be naturally represented by tensors, which are multidimensional arrays generalized from vectors and matrices. Tensor entropy is a spectral measure which can decipher topological attributes of uniform hypergraphs [11]. Here, we try to partially recover the 3D configuration of the genome using uniform hypergraphs via multi-correlation, a generalization of Pearson correlation which can measure the strength of multivariate correlation (defined in Algorithm 2 Step 3) [12]. The reconstruction is able to provide more information about genome structure and patterns, compared to the pairwise Hi-C contacts. Note that the optimal uniformity parameter $k$ depends on the internal data structure, and we leave the choice of $k$ to the users according to their data. Moreover, the constructions of adjacency tensor and degree tensor of a uniform hypergraph can be found in Chen *et al.* [11].

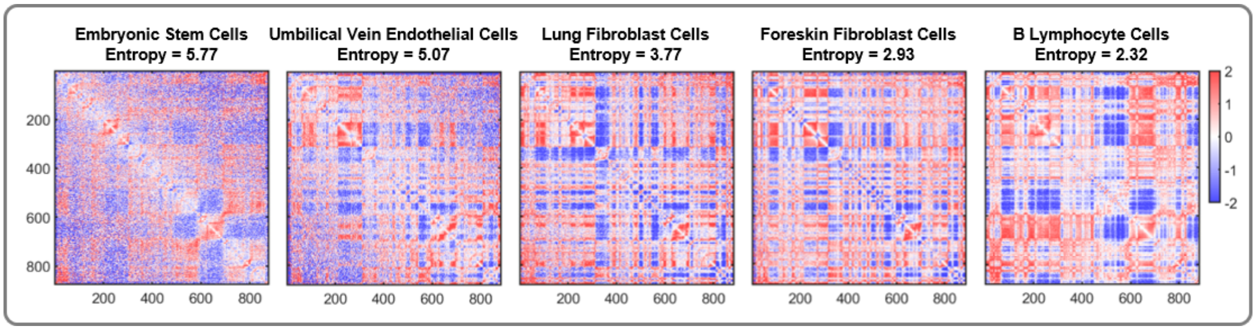


Figure S1: Entropy comparison between cell types. Hi-C matrices are shown in order of decreasing entropy (left to right). The more pluripotent cell lines (hESC, HUVEC) have a much higher entropy than the more differentiated cell lines (IMR90, HFFc6, GM12878). Intra-chromosome $\log_2$ Hi-C correlation matrices are shown at 100 kb resolution for Chromosome 14. The $y$-axis labels and color scale bar apply to all matrices in this figure.

We borrow an example of tensor entropy in the case of cellular reprogramming from Chen *et al.* [11]. The dataset contains normalized Hi-C matrices from fibroblast proliferation and MYOD1-mediated fibroblast reprogramming (MYOD1 is the transcription factor used for control) for Chromosome 14 at 1 Mb resolution with a total of 89 genomic loci. We assume $k = 3$ and $\epsilon = 0.95$ in constructing the uniform hypergraphs since third-order contacts have been shown to occur more often than pairwise contacts in the genome [13]. We can successfully detect a bifurcation in the fibroblast proliferation and reprogramming data, and accurately identify the critical transition point between cell identities during reprogramming using tensor entropy (Figure S2A). To the contrary, the network entropy cannot offer valuable insights on the bifurcation, if one analyzes the Hi-C data as adjacency matrices directly (Figure S2B). This analysis can be performed using the function tensorEntropyExample which is called by the script ExampleScript.m.

**Algorithm 2:** Tensor Entropy Computation

**Input:** Hi-C matrix $\mathbf{A} \in \mathbb{R}^{n \times n}$, uniformity parameter $k$, edge threshold $\epsilon$

**Output:** Tensor Entropy

Compute the correlation matrix $\mathbf{C} = \text{corr}(\log_2(\mathbf{A}))$

Extract all $\binom{n}{k}$ submatrices $\mathbf{R} \in \mathbb{R}^{k \times k}$ where $\mathbf{R}$ is the correlation matrix of every $k$ genomic loci

Compute the multi-correlation for every $k$ genomic loci: $\rho = (1 - \det(\mathbf{R}))^{\frac{1}{2}}$

Build a $k$-uniform hypergraph based on the multi-correlations, i.e. an edge is created if $\rho \geq \epsilon$

Compute the Laplacian tensor $\mathsf{L} = \mathsf{D} - \mathsf{A}$ where $\mathsf{D} \in \mathbb{R}^{n \times n \times \overset{k}{\cdots} \times n}$ is the degree tensor, and $\mathsf{A}$ is the adjacency tensor of the $k$-uniform hypergraph

Reshape $\mathsf{L}$ by stacking the first $k-1$ dimensions, i.e. $\mathbf{L} \in \mathbb{R}^{n^{k-1} \times n}$

Compute the singular values $\lambda_i$ of $\mathbf{L}$ using economy-size singular value decomposition ($\lambda_i$ are also called the generalized singular values of $\mathsf{L}$)

Normalize the generalized singular values: $\bar{\lambda}_j = \frac{\lambda_j}{\sum_{i=1}^{n} \lambda_i}$

Compute Tensor Entropy: **Tensor Entropy** $= -\sum_j \bar{\lambda}_j \ln \bar{\lambda}_j$

**Return:** Tensor Entropy

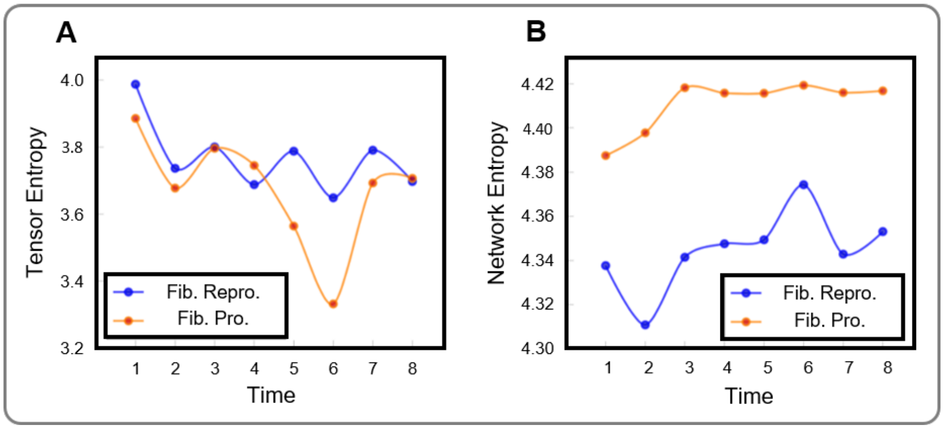


Figure S2: Cellular reprogramming features. (A) Tensor entropies of the uniform hypergraphs recovered from Hi-C measurements with multi-correlation cutoff threshold 0.95. (B) Network entropies of the binarized Hi-C matrices with weight cutoff threshold 0.95.

### LP Procedure for Comparing Hi-C Matrices

A number of methods have been developed in the Hi-C research community to compare Hi-C matrices. These methods can be broadly grouped into 2 categories: Hi-C reproducibility methods (e.g. HiCRep, HiC-spector), and Differential Chromatin Interactions (DCI) methods (e.g. SELFISH, HiCCompare, FIND, diffHic) [14, 15, 16, 17, 18, 19]. Hi-C reproducibility methods are useful for assessing the equality of Hi-C matrices genome-wide, and detecting technical bias between samples. DCI methods determine which loci have a significantly different number of Hi-C contacts between samples. All methods listed above are comparisons between only two matrices.

We pose a new method for comparing Hi-C matrices, the Larntz-Perlman procedure (LP procedure). Elements within Hi-C matrices have been shown to be normally distributed when the $\log_2$ transformation is applied to $\mathbf{A}^{(m)}$ (See Supplementary Figure S3B) [20]. For data of this form, a method for testing the equality of correlation matrices was proposed by Larntz and Perlman in 1985 [21]. We recommend that the LP procedure is performed at Hi-C resolutions $\leq$ 100 kb and that regions do not extend >5 Mb, as signal (counts) often become sparse as the genomic distance between loci increases (See Figure S3A). The full algorithm is given in Algorithm 3. The LP procedure outputs a $p$-value for the equality of the Hi-C matrices and a matrix $\mathbf{S}$ where the largest values in $\mathbf{S}$ correspond to genomic regions that are most different between all samples. This analysis can be performed using the function lpExample which is called by the script ExampleScript.m.

#### Hi-C Matrix Comparison

In order to assess the LP procedure's ability to detect differences in Hi-C matrices between samples, we have compared the LP procedure against alternate Hi-C comparison methods: HiCRep, HiC-spector, and SELFISH [14, 15, 16]. HiCRep, HiC-spector are Hi-C reproducibility metrics, while SELFISH is a DCI method that was recently shown to outperform all prior methods for DCI detection [16]. We note that there are no methods that are a direct comparison with our method. All of the alternate methods we are using here were designed to address related, but fundamentally different, questions: Hi-C reproducibility metrics for genome-wide comparisons, and DCI for loci interaction differences (not overall structural differences within a region).

There are many ways in which Hi-C matrices can be different between samples. Chromatin loops and compartments are two features that can be detected through Hi-C, and are often different between cell states [9, 22]. To assess how the LP procedure performs when these features are different between samples, we have created simulated Hi-C data sets with incremental changes in these features (See details in Supplementary

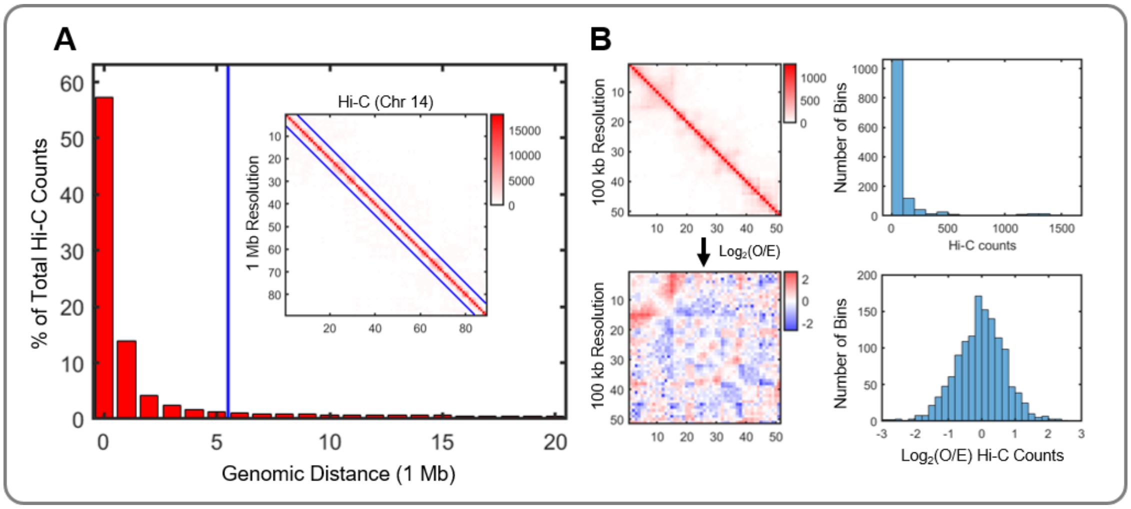


Figure S3: Hi-C normalization for the LP procedure. **(A)** Percentage of total Hi-C contacts by genomic distance. Inset figure shows a typical Hi-C intra-chromosome adjacency matrix at 1 Mb resolution. The blue line denotes where the genomic distance exceeds 5 Mb. **(B)** A 5 Mb Hi-C matrix (100 kb resolution, top left) and a histogram of the counts within the matrix (top right). When the $\log_2(O/E)$ transformation is computed on the Hi-C matrix (bottom left), elements within the matrix are normally distributed (bottom right). Here, O/E is the observed Hi-C matrix divided by the expected counts based on genomic distance.

Materials "Simulated Hi-C data"). We then determined the point at which the LP procedure, as well as alternate methods, detect differences between the matrices (See Table S1).

For Hi-C matrices with changes in loop structure as the only differences between samples, the LP procedure does not detect that the matrices are different until the loop interaction contained six times the mean number of contacts for loci at the given genomic distance. Simulated matrices for this analysis are displayed in Figure S4A. In this situation, SELFISH does the best job of detecting differences between samples. HiC-spector shows a consistent decrease in its reproducibility score, while HiCRep performs similar to the LP procedure. We note that the LP procedure, as well as HiCRep, relies on changes in the correlation between samples, and thus changes to a small number of elements within the Hi-C matrices do not affect the measurement significantly.

For Hi-C matrices with changes in the compartment structure of a single bin (genomic region) as the only differences between samples, the LP procedure performs very well. Simulated matrices for this analysis are displayed in Figure S4B. SELFISH detects differences between all samples, but also detects differences when only a small amount of noise is added to the original matrix. The LP procedure is robust to noise and shows a trend consistent with the amount of change created. The HiC-spector reproducibility measurement shows no clear trend as the matrices diverge, and the LP procedure detects more subtle changes relative to HiCRep. Compartment changes are often observed in time series Hi-C matrices as cells transition between cell states, either through reprogramming or differentiation [23].

### Simulated Hi-C Data

Simulated Hi-C data was created to compare the LP procedure against alternative Hi-C comparison methods. Two distinct simulated data sets were created to have: (1) changes in chromatin loop structure and (2) changes in chromatin compartment structure. Chromatin loops and chromatin compartments are two features that have been used to characterize Hi-C structure. Both simulated data sets are created by perturbing data from real Hi-C matrices: a 400 kb region (10 kb resolution) is used for the chromatin loop data set and a 2 Mb region (50 kb resolution) is used for the chromatin compartment data set.

One additional simulated matrix was also created by adding a small amount of random noise to the 400

**Algorithm 3:** LP procedure

**Input:** Hi-C matrices $\mathbf{A}^{(m)} \in \mathbb{R}^{n \times n}$, $m = 1, \ldots, T$
**Output:** $p$-value $p$, and test statistic $\mathbf{S}$

1 Compute the correlation matrix $\mathbf{C}^{(m)} = \text{corr}(\log_2(\mathbf{A}^{(m)}))$. Define the corresponding population correlation matrices as $\mathbf{P}^{(m)}$

2 Define the null hypothesis

$$H_0 : \mathbf{P}^{(1)} = \cdots = \mathbf{P}^{(T)}$$

3 Compute the Fisher z-transformation $\mathbf{Z}^{(m)}$. Elements in $\mathbf{Z}^{(m)}$ and $\mathbf{C}^{(m)}$ are denoted as $z_{ij}^{(m)}$ and $c_{ij}^{(m)}$, respectively, and

$$z_{ij}^{(m)} = \frac{1}{2}\ln\left[\frac{1 + c_{ij}^{(m)}}{1 - c_{ij}^{(m)}}\right]$$

4 Form $\mathbf{S}$, where elements in $\mathbf{S}$ are denoted as $s_{ij}$ and

$$s_{ij} = (n-3)\sum_{m=1}^{T}(z_{ij}^{(m)} - \bar{z}_{ij})^2, \;\; \bar{z}_{ij} = T^{-1}\sum_{m=1}^{T} z_{ij}^{(m)}$$

5 Calculate the test statistic

$$T = \max_{1 \le i < j \le n} s_{ij}$$

6 Reject $H_0$ at level $\alpha$ if $T > \chi^2_{T-1,\epsilon(\alpha)}$, where $\chi^2_{T-1,\epsilon(\alpha)}$ is the chi-squared distribution with $T-1$ degrees of freedom and $\epsilon(\alpha) = (1-\alpha)^{2/n(n-1)}$ is the Šidák correction

7 Calculate $p$, the level $\alpha$ at which $T > \chi^2_{T-1,\epsilon(\alpha)}$

**Return:** $p$ and $\mathbf{S}$

kb region (10 kb resolution) Hi-C matrix. A random number sampled from a normal distribution, $\mathcal{N}(0, 0.05)$, is added to each element in $\log_2(\mathbf{A})$. This matrix is used to determine how robust Hi-C matrix comparison methods are to small amounts of noise.

For each simulated data set, 10 matrices were created that are incrementally more divergent from the original Hi-C matrix. For changes in loop structure, counts were added to a specific off-diagonal region, following a 2D Gaussian distribution ($\sigma = 1$), to model a chromatin loop structure. For changes in compartment structure, counts aligned to a specific genomic region (bin) were changed to decrease the correlation coefficient between the specified bin in the simulated $\log_2$ Hi-C matrix and the specified bin in the original $\log_2$ Hi-C matrix. These changes reflect what would be observed if the specified bin was changing its compartment structure. Methods to recreate the simulated Hi-C matrices are provided within the 4DNvestigator.

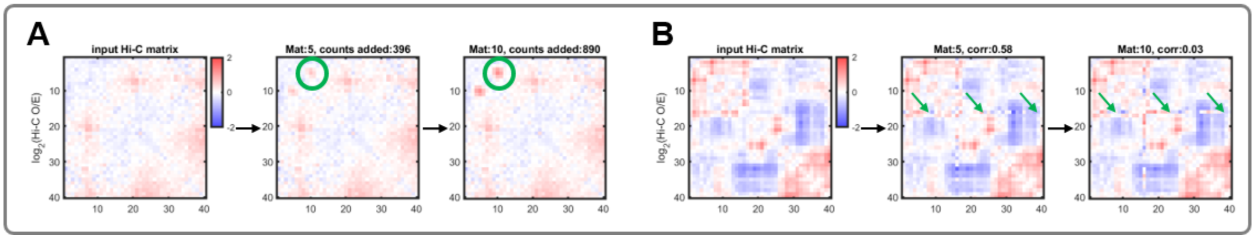


Figure S4: Simulated Hi-C data. **(A)** Simulated Hi-C matrices for loop changes (See Supplementary Table S2). The original Hi-C matrix is shown on the left, with matrices that are increasingly more divergent from left to right. Green circles indicate where the matrix has been perturbed to change the chromatin loop structure. **(B)** Simulated Hi-C matrices for compartment changes (See Supplementary Table S2). The original Hi-C matrix is shown on the left, with matrices that are increasingly more divergent from left to right. Green arrows indicate where the matrix has been perturbed to change the chromatin compartment structure.

| Method | Counts added 53 | 160 | 266 | 372 | 479 | Correlation 1 (noise) | 0.87 | 0.57 | 0.2 | -0.26 | -0.65 |
|---|---|---|---|---|---|---|---|---|---|---|---|
| LP ($p$-value) | 1.0 | 1.0 | 0.99 | 0.41 | 0.045 | 1.0 | 1.0 | 2.1E-03 | 2.5E-07 | 1.7E-13 | 0.0 |
| SELFISH ($p$-value) | 0.0 | 0.0 | 0.0 | 0.0 | 0.0 | 0.02 | 2.9E-05 | 1.4E-08 | 1.3E-10 | 3.5E-13 | 2.1E-07 |
| HiCRep (SCC) | 1.0 | 1.0 | 1.0 | 1.0 | 1.0 | 1.0 | 1.0 | 0.99 | 0.98 | 0.97 | 0.96 |
| HiC-spector (Q) | 0.99 | 0.98 | 0.96 | 0.94 | 0.92 | 0.39 | 0.43 | 0.33 | 0.48 | 0.44 | 0.50 |

Table S2: Hi-C matrix comparison methods. "Counts added" refers to the number of counts added to the simulated Hi-C matrix at the specified location. "Correlation" refers to the correlation between the selected column in the original Hi-C matrix, and the same column in the simulated Hi-C matrix. We note here that each method outputs a different unit, as specified within the table. The lowest $p$-value output from SELFISH is given here.

### References


[1] Belkin, M. and Niyogi, P. (2001) Laplacian Eigenmaps and Spectral Techniques for Embedding and Clustering. *Nips,* **14**, 585–591.

[2] Van Der Maaten, L. and Hinton, G. E. (2008) Visualizing high-dimensional data using t-sne. *Journal of Machine Learning Research,* **9**, 2579–2605.

[3] McInnes, L., Healy, J., and Melville, J. (2018) Umap: Uniform manifold approximation and projection for dimension reduction. *arXiv preprint arXiv:1802.03426,*.

[4] Cover, T. M. and Thomas, J. A. (2012) Elements of information theory, John Wiley & Sons, .

[5] Macarthur, B. D. and Lemischka, I. R. (2013) Statistical mechanics of pluripotency. *Cell,* **154**(3), 484–489.

[6] Rajapakse, I., Groudine, M., and Mesbahi, M. (2012) What can systems theory of networks offer to biology?. *PLoS computational biology,* **8**(6), e1002543.

[7] Meshorer, E. and Misteli, T. (2006) Chromatin in pluripotent embryonic stem cells and differentiation. *Nature reviews Molecular cell biology,* **7**(7), 540.

[8] Dekker, J., Belmont, A. S., Guttman, M., Leshyk, V. O., Lis, J. T., Lomvardas, S., Mirny, L. A., O'Shea, C. C., Park, P. J., Ren, B., et al. (2017) The 4D nucleome project. *Nature,* **549**(7671), 219–226.

[9] Rao, S. S., Huntley, M. H., Durand, N. C., Stamenova, E. K., Bochkov, I. D., Robinson, J. T., Sanborn, A. L., Machol, I., Omer, A. D., Lander, E. S., et al. (2014) A 3D Map of the Human Genome at Kilobase Resolution Reveals Principles of Chromatin Looping. *Cell,* **159**(7), 1665–1680.

[10] Darrow, E. M., Huntley, M. H., Dudchenko, O., Stamenova, E. K., Durand, N. C., Sun, Z., Huang, S.-C., Sanborn, A. L., Machol, I., Shamim, M., et al. (2016) Deletion of DXZ4 on the human inactive X chromosome alters higher-order genome architecture. *Proceedings of the National Academy of Sciences,* **113**(31), E4504–E4512.

[11] Chen, C. and Rajapakse, I. (2020) Tensor Entropy for Uniform Hypergraphs. *IEEE Transactions on Network Science and Engineering,* **7**(4), 2889–2900.

[12] Wang, J. and Zheng, N. (2014) Measures of Correlation for Multiple Variables. *arXiv preprint arXiv:1401.4827,*.

[13] Ulahannan, N., Pendleton, M., Deshpande, A., Schwenk, S., Behr, J. M., Dai, X., Tyer, C., Rughani, P., Kudman, S., Adney, E., et al. (2019) Nanopore sequencing of DNA concatemers reveals higher-order features of chromatin structure. *bioRxiv,* p. 833590.

[14] Yang, T., Zhang, F., Yardimci, G. G., Song, F., Hardison, R. C., Noble, W. S., Yue, F., and Li, Q. (2017) HiCRep: assessing the reproducibility of Hi-C data using a stratum- adjusted correlation coefficient. *Genome Research,* **27**(11), 1939–1949.

[15] Yan, K. K., Yardlmcl, G. G., Yan, C., Noble, W. S., and Gerstein, M. (2017) HiC-spector: A matrix library for spectral and reproducibility analysis of Hi-C contact maps. *Bioinformatics,* **33**(14), 2199–2201.

[16] Ardakany, A. R., Ay, F., and Lonardi, S. (2019) Selfish: Discovery of Differential Chromatin Interactions via a Self-Similarity Measure. *bioRxiv,* p. 540708.

[17] Djekidel, M. N., Chen, Y., and Zhang, M. Q. (2018) FIND: DifFerential chromatin INteractions Detection using a spatial Poisson process. *Genome Research,* **28**(3), 412–422.

[18] Stansfield, J. C., Cresswell, K. G., Vladimirov, V. I., and Dozmorov, M. G. (2018) HiCcompare: An R-package for joint normalization and comparison of HI-C datasets. *BMC Bioinformatics,* **19**(1), 13–16.

[19] Lun, A. T. and Smyth, G. K. (2015) diffHic: A Bioconductor package to detect differential genomic interactions in Hi-C data. *BMC Bioinformatics,* **16**(1), 1–11.

[20] Calandrelli, R., Wu, Q., Guan, J., and Zhong, S. (2018) GITAR: An Open Source Tool for Analysis and Visualization of Hi-C Data. *Genomics, Proteomics and Bioinformatics,* **16**(5), 365–372.

[21] Larntz, K. and Perlman, M. D. (1985) A simple test for the equality of correlation matrices. *Rapport technique, Department of Statistics, University of Washington,* **141**.

[22] Lieberman-aiden, E., Berkum, N. L. V., Williams, L., Imakaev, M., Ragoczy, T., Telling, A., Amit, I., Lajoie, B. R., Sabo, P. J., Dorschner, M. O., et al. (2009) Comprehensive Mapping of Long-Range Interactions Reveals Folding Principles of the Human Genome. *Science,* **326**, 289–294.

[23] Dixon, J. R., Jung, I., Selvaraj, S., Shen, Y., Antosiewicz-Bourget, J. E., Lee, A. Y., Ye, Z., Kim, A., Rajagopal, N., Xie, W., et al. (2, 2015) Chromatin architecture reorganization during stem cell differentiation. *Nature,* **518**(7539), 331–336.
